# Precise measurement of nanoscopic septin ring structures in deep learning-assisted quantitative superresolution microscopy

**DOI:** 10.1101/2021.12.28.474382

**Authors:** Amin Zehtabian, Paul Markus Müller, Maximilian Goisser, Leon Obendorf, Lea Jänisch, Nadja Hümpfer, Jakob Rentsch, Helge Ewers

## Abstract

The combination of image analysis and fluorescence superresolution microscopy methods allows for unprecedented insight into the organization of macromolecular assemblies in cells. Advances in deep learning-based object recognition enables the automated processing of large amounts of data, resulting in high accuracy through averaging. However, while the analysis of highly symmetric structures of constant size allows for a resolution approaching the dimensions of structural biology, deep learning methods are prone to different forms of bias. A biased recognition of structures may prohibit the development of readouts for processes that involve significant changes in size or shape of amorphous macromolecular complexes. What is required to overcome this problem is a detailed investigation of potential sources of bias and the rigorous testing of trained models using real or simulated data covering a wide dynamic range of possible results. Here we combine single molecule localization-based superresolution microscopy of septin ring structures with the training of several different deep learning models for a quantitative investigation of bias resulting from different training approaches and finally quantitative changes in septin ring structures. We find that trade-off exists between measurement accuracy and the dynamic range of recognized phenotypes. Using our trained models, we furthermore find that septin ring size can be explained by the number of subunits they are assembled from alone. Our work provides a new experimental system for the investigation of septin polymerization.

## Introduction

Light microscopy is a technique of fundamental importance in cell biology and with the development of superresolution techniques has allowed unprecedented insight into cellular processes on the brink of structural biology^1–7^. Machine learning-based computational image processing furthermore allows for high-throughput analysis of even complex phenotypes in cells^8–12^ and has accelerated progress in cell biological discovery. In recent years a number of open source deep learning platforms have been made available^13–22^ that require minimal know-how on the user side. At the same time, methods have been developed to reduce background^23,24^ and to accelerate image processing in superresolution microscopy^25–27^. Especially in superresolution microscopy, image processing allows for ultrahigh resolution of multiprotein complexes such as nuclear pores^3,28^ or centrioles^29,30^. However, these are highly symmetric structures of constant size that bear several symmetry axes. As a result, they contain intrinsic means to allow for accurate averaging. For superresolution microscopy and averaging to harness its full power with the help of deep learning-based image processing to improve resolution of more amorphous structures and to develop quantitative readouts for structural changes it remains to be determined to what extent deep learning methods can handle changes in organization or even size of target structures. Overfitting may make it impossible to detect even small changes in morphology or size, on the other hand poor fitting may not allow for accurate measurements.

Here we combine deep learning-assisted image analysis based on an open-source platform with single molecule localization-based superresolution microscopy (SMLM) of subresolution septin ring structures. Septins are the fourth cytoskeleton^31,32^ and assemble from nonpolar heteromultimeric rod-shaped complexes into ring-like structures of subresolution size when not associated to tubulin or actin^33^. These rings are free of actin and are thought to represent *bona fide* septin filamentous polymers^34^. The composition of septin complexes plays an important role in septin function^35,36^ and septin ring size depends on the composition of septin complexes^37^. Septin ring size may thus be a long sought-after readout for complex assembly and composition. Here we ask what factors in supervised deep learning affect ring recognition and finally, accuracy of ring size measurements in native and perturbed cells with differing ring size. To do so, we train six different models on data annotated according to several different parameters and by three different experts and validate them on cellular and synthetic SMLM datasets. We find that while most models readily recognize ring-structures in the size range expected in native cells and allow for highly accurate determination of septin ring diameter from SMLM data, only few models allow for the accurate recognition and measurement of rings of differing size. We furthermore show that septin ring size can be explained by septin complex composition alone, and we thus provide a new experimental paradigm for the investigation of septin complex assembly. Our results demonstrate that the combination of deep learning with SMLM can provide accurate readouts of changes in the size of amorphous multiprotein structures and that experimental design and careful validation are essential for the generation of reliable experimental pipelines.

## Results

We chose septin rings as a model to test the robustness and accuracy of deep learning-assisted superresolution microscopy for measurements of subresolution cellular structures. The septins are a family of GTP-binding proteins that comprise the fourth cytoskeleton. Septins can assemble into filaments themselves or attach to actin stress fibers, however, when actin stress fibers are collapsed via the use of cytochalasin D, they form ring structures of around 350 nm diameter^33,37^ (Figure 1A). When imaged in SMLM, rings appear as homogenously sized, symmetric rings with a thin perimeter (Figure 1B), but also incomplete rings, arcs, small filamentous structures and aggregates can be found. When we acquired many cells in SMLM imaging after Cytochalasin D treatment and selected hundreds of rings from the stained septin structures, we could average them for accurate determination of their diameter of 343.9 ± 5.0 nm (n = 181, Figure 1C). We concluded that we could accurately determine septin ring diameter using this method.

**FIGURE 1:**
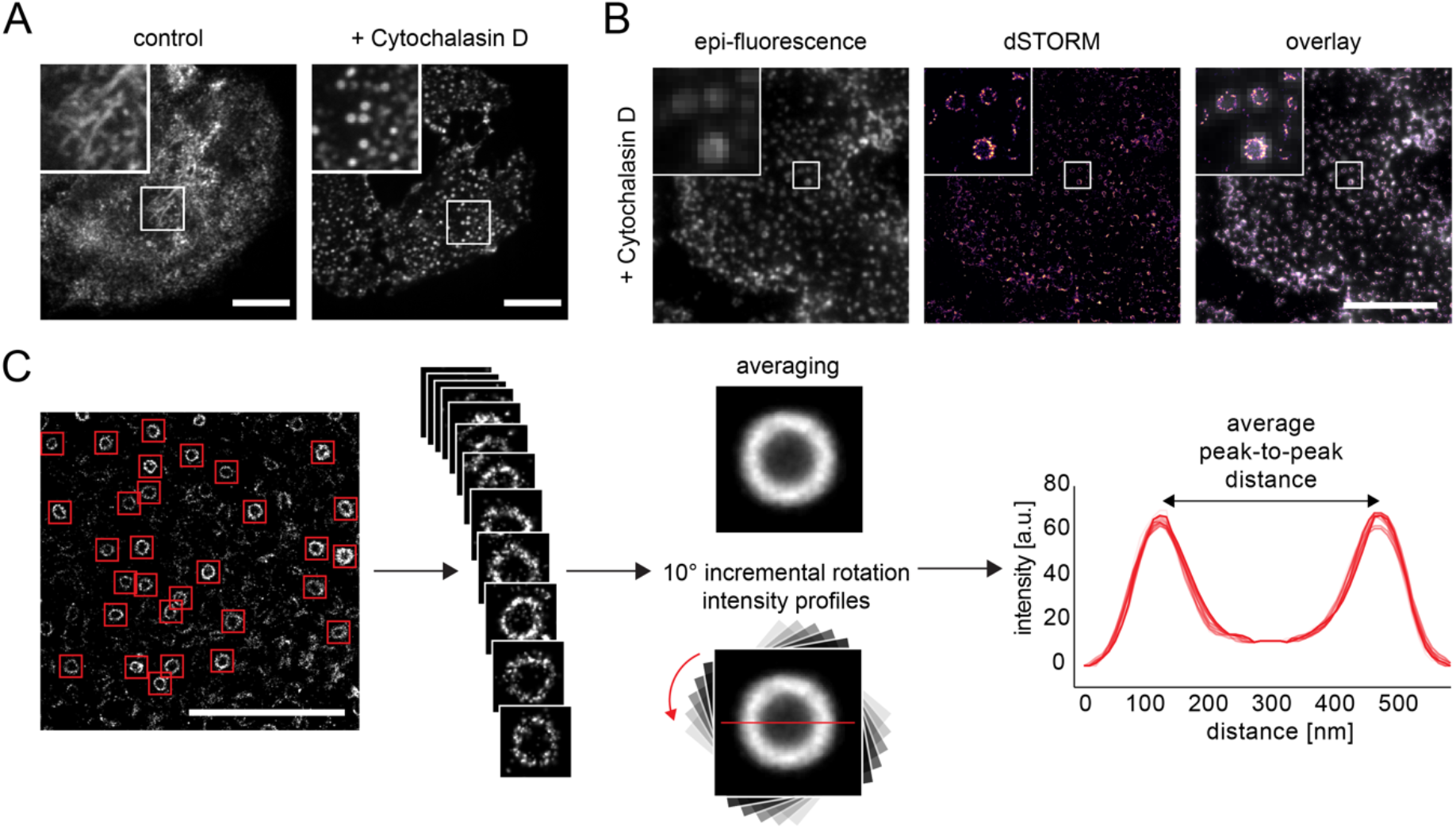
Superresolution microscopy assay for quantification of septin ring diameter. **(A)** Genome-edited NRK52E cells expressing mEGFP-Septin-2 from both alleles were treated with 5 μM Cytochalasin D for 30 min (or DMSO as control) prior to fixation, resulting in the formation of ring-like structures. Images of mEGFP fluorescence were taken by spinning disc confocal microscopy. Scale bar: 10 μm. **(B)** Septin rings can be readily resolved by single molecule localization-based superresolution microscopy. NRK52E cells were treated with 5 μM Cytochalasin D for 30 min and immunostained against Septin-7. Images were taken by conventional epi-fluorescence microscopy and by dSTORM. Scale bar: 10 μm. **(C)** Scheme of the procedure for the measurement of septin ring diameter. Left to right: Septin rings from superresolution microscopy images were cropped and averaged. The intensity profile of the averaged rings was measured at 18 successive 10° incremental rotations and the peak-to-peak distance of all of these profiles was averaged to return the ring diameter. Scale bar: 5 μm.

We next aimed to use a widely available and versatile deep learning network to train it to recognize septin rings. We decided to use the ZeroCostDL4Mic^13^ platform which provides free cloud-based computation and the StarDist^38^ network that has been widely used for detection of cellular structures. However, since it had not been used for ring detection we decided to test, if it could be trained from scratch to recognize septin rings in SMLM reconstructions. To do so, we used an annotated dataset of 496 images for training and adjusted parameters of the StarDist network until we could reliably reach convergence of our model.

We next aimed to ask, how annotation bias may affect model learning and the capability of the model to detect phenotypes strongly different from the wildtype *(wt)* situation in our images. In cellular SMLM images, septin rings are of variable circularity and completeness in labeling and occasionally touch. We hence decided to test if higher or lower stringency in determining a ring would be suitable for training and if masks precisely following the ring outline would lead to more accurate results. After training, we furthermore asked, whether the deep learning models would be able to accurately recognize and measure also significantly smaller or bigger rings.

For training, we generated a large amount of SMLM images of cytochalasin D-treated cells, which we separated into a training set of 496 images and a cellular holdout test dataset of 32 images. We then designed a framework to train deep learning models using six different annotation strategies and to test the models on the cellular holdout data, the synthetic data and a biological test case, that is rings devoid of Septin-9 (Figure 2). We asked one human expert to annotate in three different ways in terms of accuracy of the masks (how does it fit to the ring) and labeling stringency (how circular and/or continuous do rings have to be) and after that asked two more human experts to individually annotate with custom masks and low stringency. A final sixth model was trained using the consensus masks. All trained models resulted in robust recognition of rings in the cellular test dataset of 32 SMLM images and we benchmarked the models in terms of precision (fraction of recognized objects that are present in ground truth), recall (fraction of objects in ground truth that are recognized) and measured average ring size compared to ground truth (Figure 2). We then went on to run the models on synthetic SMLM data generated using FluoSim^39^ in which ring sizes from 125 to 500 nm were simulated. Finally, we tested the two best performing models on a cellular example of ring size change, cells in which Septin-9 had been removed from the complex, leading to significantly smaller rings^37^.

**FIGURE 2:**
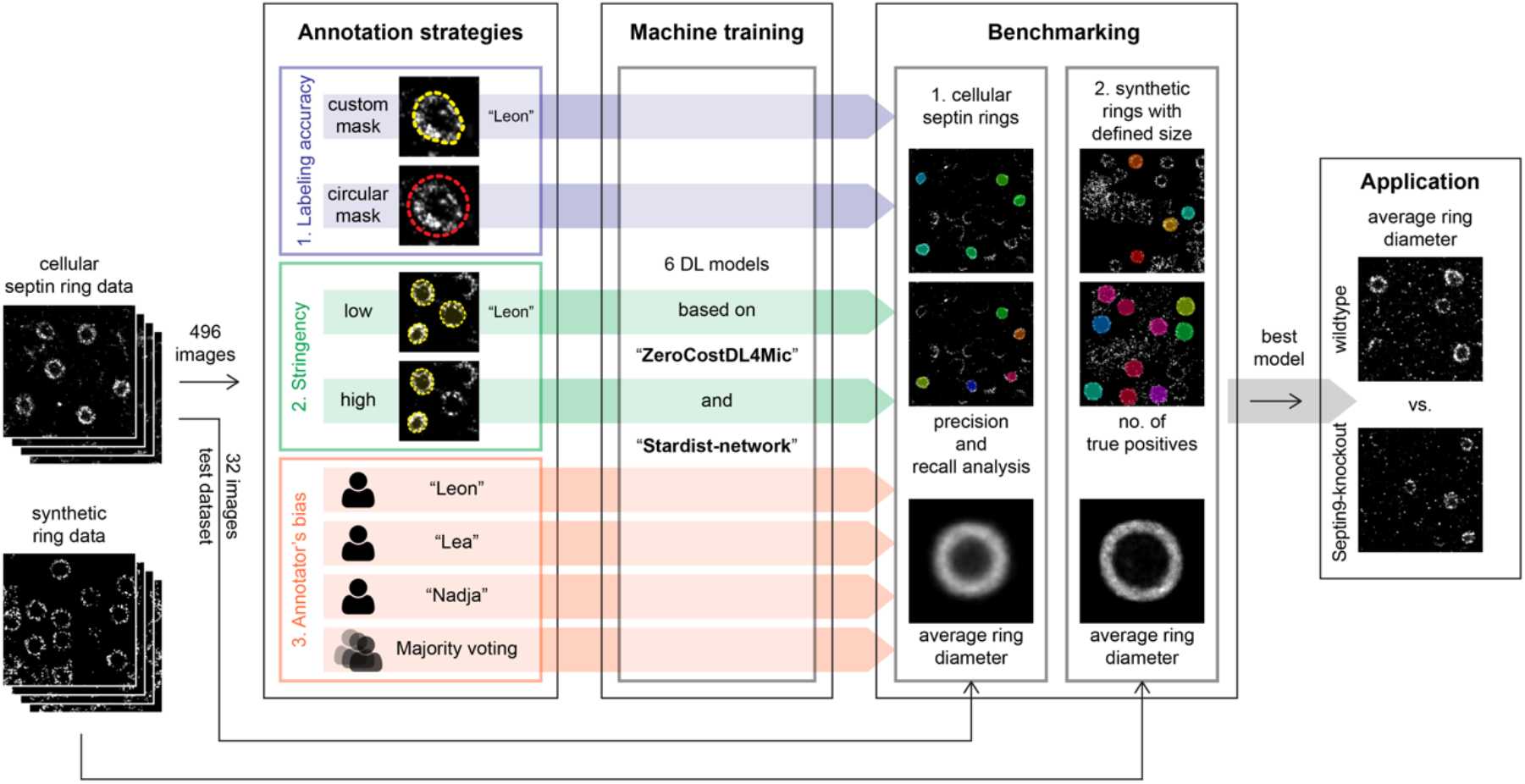
General annotation, training and benchmarking strategy for deep learning-assisted septin ring recognition and measurement. 528 dSTORM images of septin rings in NRK52E cells were generated and used for annotation and training (496 images) or as holdout test dataset (32 ground truth images). To benchmark annotation strategies for deep learning, images were annotated with different labeling accuracy, annotation stringency, and by three independent human experts as well as the majority voting thereof. For each annotation strategy we generated a deep learning models based on the ZeroCostDL4Mic platform using the StarDist network. For benchmarking, we compared the ability of the models to recognize the rings and to return the correct ring diameter, both in cellular data and synthetic data with rings of known diameter. The best performing models were then applied to measure the diameter of septin rings in knockout cells.

To evaluate the influence of labeling strategies on recognition by the deep learning models, a septin biologist recognized septin rings in the cellular test dataset of SMLM data. These we accepted as *bona fide* septin rings and the resulting dataset and average diameter are in the following termed ground truth. We then asked a human expert, “Leon”, to annotate the training set according to three different strategies. The first strategy used low stringency in recognition as a septin ring and required the expert to draw a custom mask around each ring (Figure 2). The second strategy required the expert to use low stringency in septin ring recognition, but to use a circular mask or, to be more precise, a convex circular-shape mask with flexible size that covered the target ring for annotation. The third strategy required the expert to apply high stringency in selecting rings and to use a custom drawn mask. When the annotation-specific models were trained and applied to the cellular test dataset, we found that all models recognized a large fraction of rings reliably when compared to ground truth albeit with varying precision and recall (Figure 3A). Specifically, the low stringency, custom mask trained model led to the highest recall with 89.5% of rings found that were present in the ground truth. On the other hand, the high stringency, custom mask trained model led to the highest precision with 89% of all recognized rings being true positives, but recall was low. The circular mask, low stringency model was of intermediate precision and recall and no model showed precision nor recall below 50%. Overall, we concluded that our models could reliably recognize septin rings in our cells.

**FIGURE 3:**
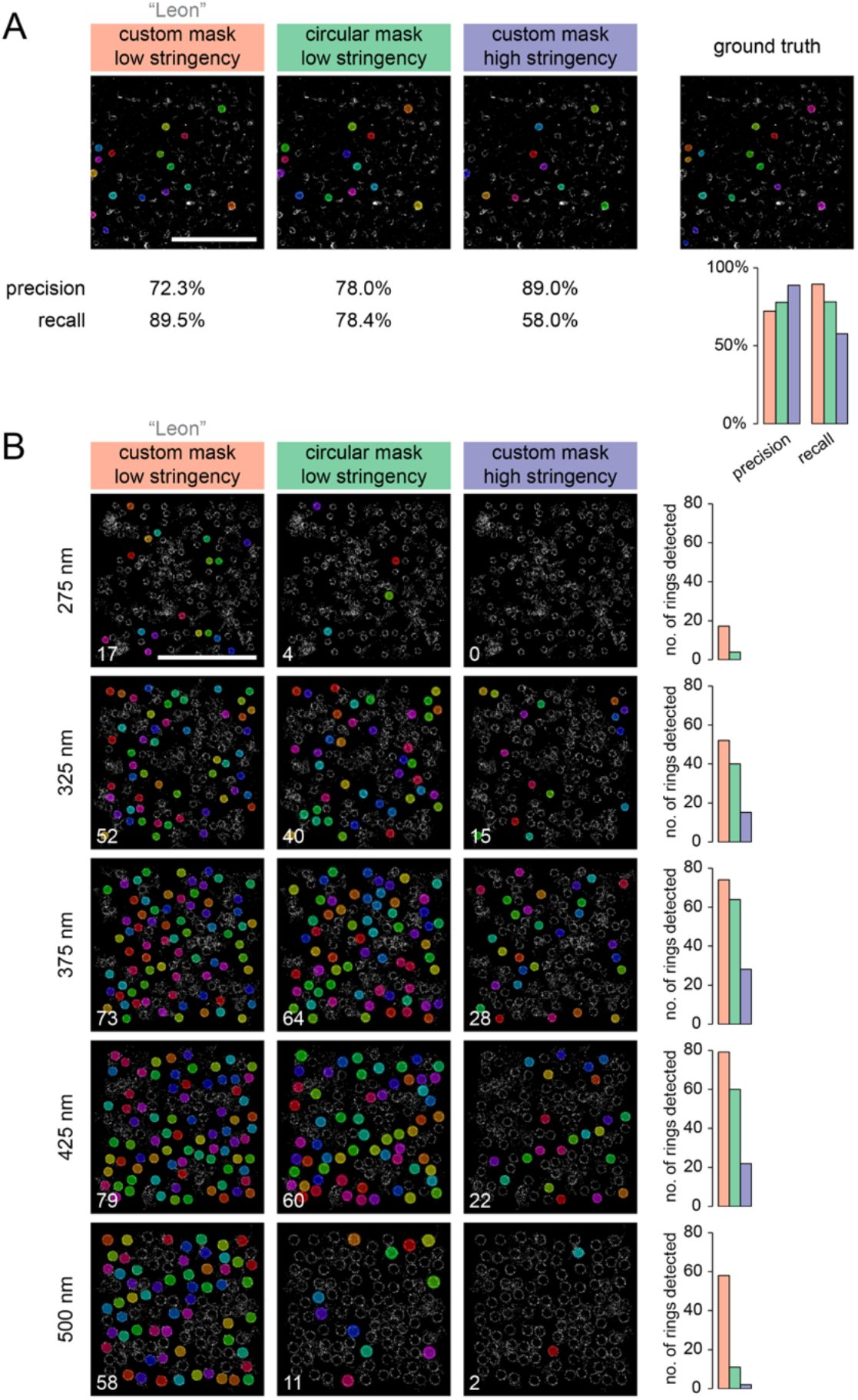
Effect of annotation accuracy and stringency on the ability of deep learning models to detect rings of different size. **(A)** Overlay of randomly colored masks of recognized septin ring structures in dSTORM images by 3 different deep learning models: custom mask and low stringency model (aka “Leon”); circular mask and low stringency model; custom mask and high stringency model. The rings detected in a representative dSTORM reconstruction from the holdout data are shown together with the respective ground truth. Precision and recall calculations are shown below. Scale bar: 5 μm. **(B)** Overlay of randomly colored masks of recognized ring structures in synthetic dSTORM images with predetermined ring diameters from 275 to 500 nm as recognized by the deep learning models as in (A). Number of detected rings in each condition is shown on the bottom left of each image and plotted on the right. Scale bar: 5 μm.

When the composition of septin complexes is changed, the diameter of septin rings changes as well^37^, suggesting that ring size may provide a readout for septin complex assembly and composition. However, to accurately measure the diameter of rings with different sizes, our models must recognize rings of even strongly differing diameters for an accurate readout of ring-size changes. We thus aimed to test whether one of our training strategies would lead to a bias in the recognition of differently sized rings and only recognize rings in the size-range found in our untreated cells. To do so, we analyzed a panel of synthetic SMLM test datasets in which rings of a predefined diameter from 275 to 500 nm, respectively, were present. When we then applied our models to recognize rings in these synthetic test datasets, we found striking differences in ring recognition (Figure 3B). While the low stringency, custom mask model (hereafter called “Leon”) reliably recognized rings in all test cases, the other two models failed to reliably recognize rings for cases significantly below or above the ring sizes present in our cells. We concluded that the custom mask, high stringency model (hereafter called “high stringency”) and the circular mask, low stringency model (hereafter called “circular mask”) were biased for rings in the size range found in cells and that the training strategy significantly impacts recognition of septin rings towards superresolution measurements.

Next, we aimed to investigate how three different experts using the same annotation strategy would influence ring recognition by the resulting models. We asked two additional experts, “Lea” and “Nadja” to annotate the cellular SMLM training dataset according to the low stringency, custom mask strategy. From the annotations of the three experts “Leon”, “Lea” and “Nadja” we furthermore created a “majority voting” annotation. When we then trained additional models according to the “Lea”, “Nadja” and “majority voting” annotations on our cellular test dataset, we found that all models exhibited a precision above 60% and a recall above 80% when compared to ground truth (Figure 4A). The “Nadja” model exhibited an exceptional recall of 92.8%, but only 60.4% precision, meaning that the “Nadja” model recognized many objects that were not found in the ground truth. The performance of the “majority voting” model was very similar to that of the “Leon” model. We concluded that indeed the specific annotation style of different experts influences ring recognition significantly. When we next applied the “Lea”, “Nadja” and “majority voting” models to the panel of synthetic test datasets and compared them to the “Leon” model, we found significant discrepancies in performance (Figure 4B).

**FIGURE 4:**
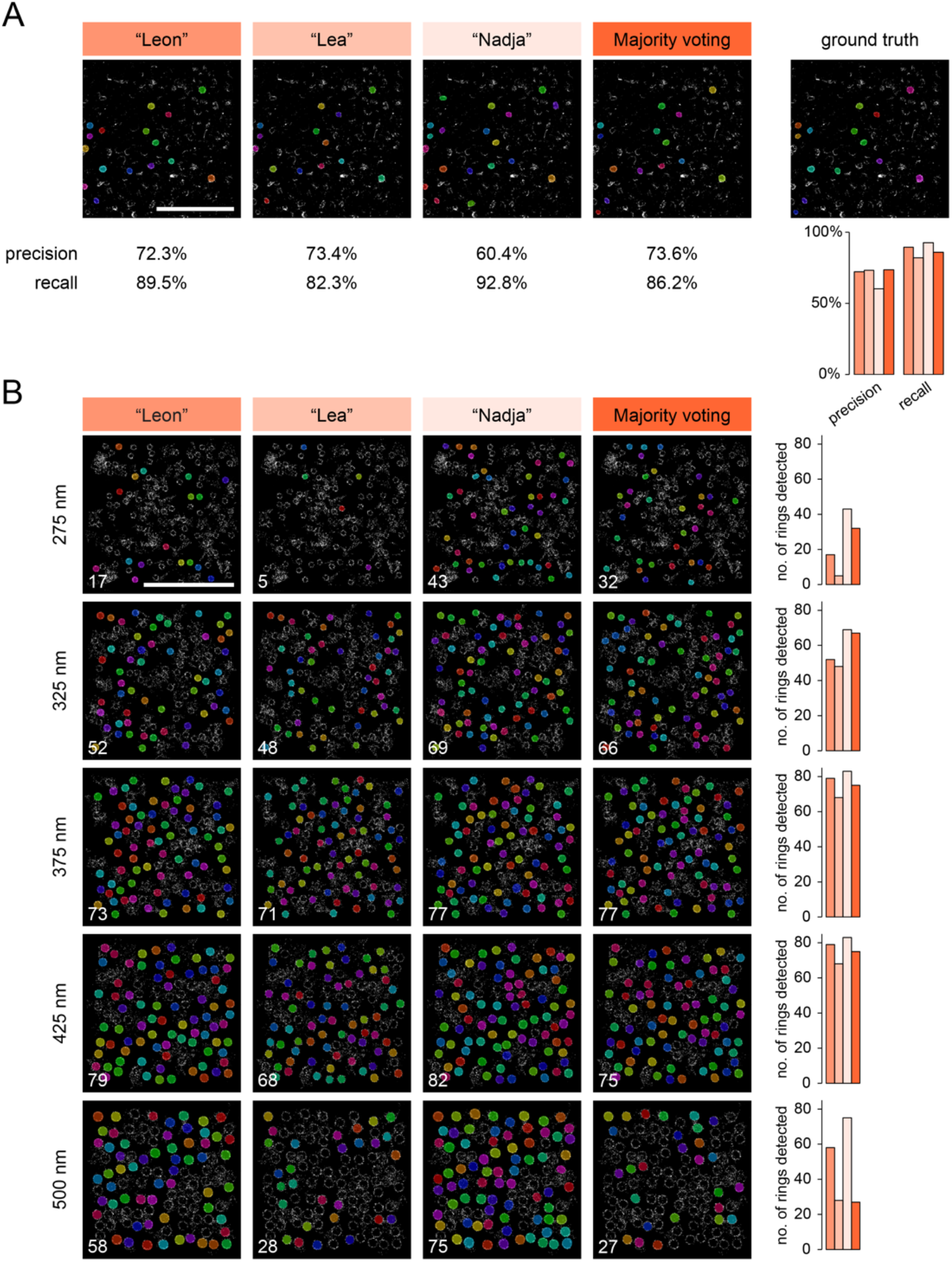
Effect of annotator’s bias on the ability of deep learning models to recognize rings of different size. **(A)** Overlay of randomly colored masks of recognized septin ring structures in dSTORM images by 4 different deep learning models: custom mask and low stringency model each for individual annotators “Leon”, “Lea”, “Nadja” and for majority voting model. The rings detected in a representative dSTORM reconstruction from the holdout data are shown together with the respective ground truth. Precision and recall calculations are shown below. Scale bar: 5 μm. **(B)** Overlay of randomly colored masks of recognized ring structures in synthetic dSTORM images with predetermined ring diameters from 275 to 500 nm as recognized by the deep learning models as in (A). Number of detected rings in each condition is shown on the bottom left of each image and plotted on the right. Scale bar: 5 μm.

Both the “Nadja” and the “majority voting” models reliably outperformed the other models for smaller rings, while for rings in the range of those found in cells, all models performed equally well. We also tested rings below 275 nm in size (data not shown). For the smallest size of 125 nm, no model could recognize any rings in synthetic SMLM test datasets. Only the “Nadja” model could recognize rings reliably from a diameter of 245 nm upward, making it clearly stand out from the other models. Large (500 nm) rings on the other hand were readily recognized by the “Nadja” and “Leon” models, while the “Lea” and “majority voting” models recognized less than half of the rings detected by either the “Nadja” or “Leon” models. We concluded that the bias introduced by individual annotators has significant consequences on the recognition of septin rings in SMLM data.

We next aimed to ask, to what extent these biases in septin ring annotation would impact the accurate measurement of septin ring diameter in datasets analyzed by our models. To do so, we averaged the recognized septin rings for every model and condition and measured the diameter in nm as described in Figure 1C. When we compared the results of our models on the cellular SMLM test dataset, we found that all models resulted in a measurement with an accuracy within 3% of the 343.9 nm found in the ground truth (Figure 5A). We next aimed to ask, if we could measure differences as small as 5 nm in ring diameter. To do so, we generated synthetic SMLM test datasets with diameters in 5 nm steps from 350 nm, to 380 nm. When we ran our models on these datasets and measured the average diameters of the respectively recognized rings, we found that all models resulted in very accurate measurements of the respective ring sizes, with errors maximally 3 - 4 nm (Figure 5B). Overall, the highest spread was found for the “circular mask” and “high stringency” models (Figure 5C), while the residuals were lowest for the “majority voting” model (Figure 5D). We concluded that all models were suitable to automatically extract and measure ring size in native cellular SMLM datasets. When we plotted the measured ring diameter against ground truth ring diameter in synthetic SMLM test data, we found that all models exhibited excellent linearity over the range from 275 to 500 nm with R^2^-values above 0.94 (Figure 6A). While thus all models were very precise in their measurement of ring diameter, a clear dependence of measurement accuracy on the number of recognized rings was found (Figure 6B), suggesting that the generation of higher amounts of data could further ameliorate measurement accuracy for the less well performing models. In Figure 6C, we summarize the data from Figures 3–5 in an accessible overview according to the z-scores for ring recognition (top) and ring diameter measurement (bottom). Found values are horizontally z-scored, thus emphasizing differences between models even if they may be small in absolute size and describing relative performance. However, this overview allows for general observations.

**FIGURE 5:**
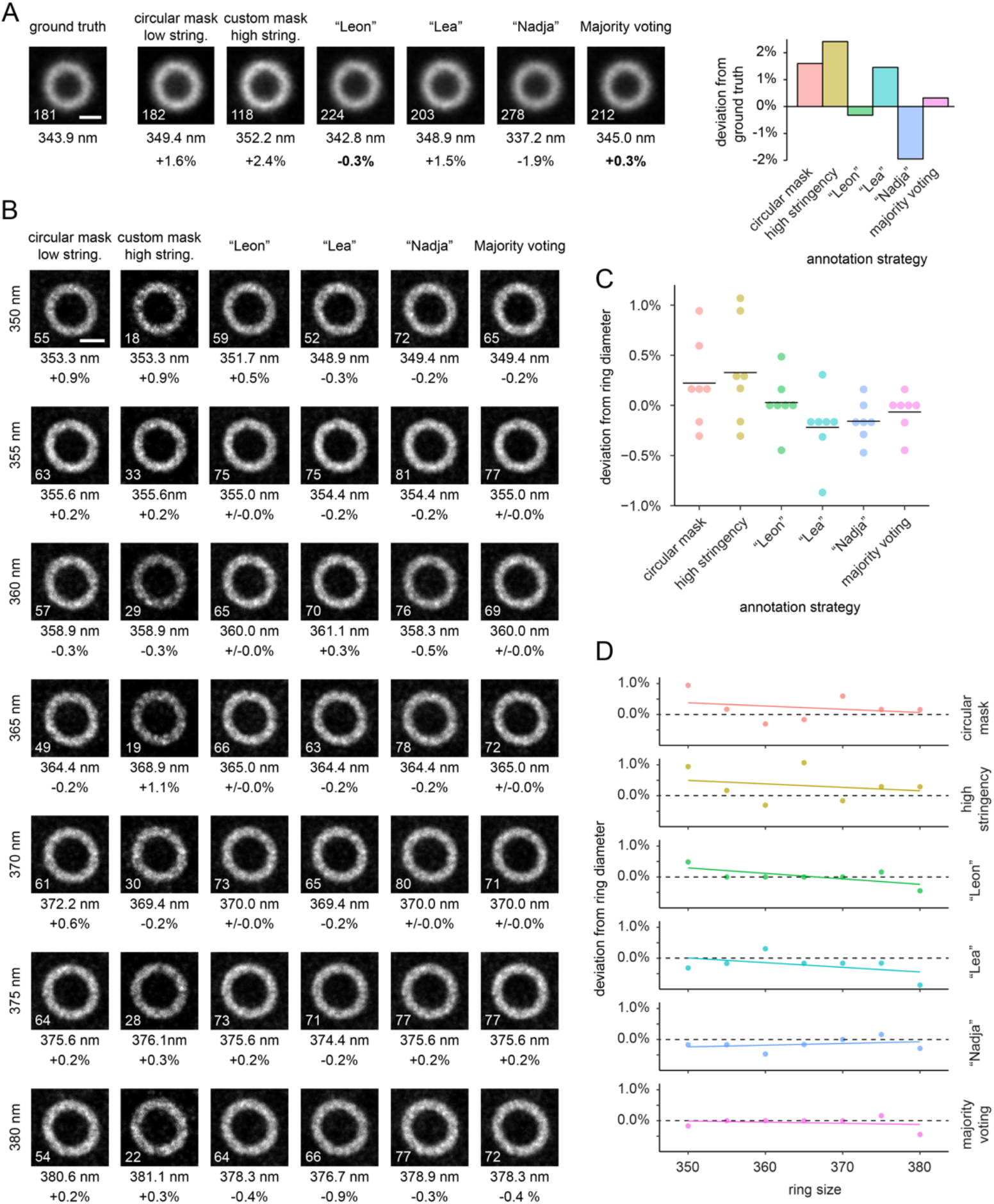
Quantification of the precision of deep learning-based ring measurements. **(A)** Results for septin ring diameter measurements in cellular dSTORM holdout data by the respective deep learning models. Averaged recognized ring images are shown for each model and average ring diameter and deviation from ground truth are shown below the images. Number of rings averaged is shown on the bottom left of each image. The plot on the right shows the deviation of measured septin ring diameter from ground truth for each model in %. Scale bar: 200 nm. **(B)** Results for septin ring diameter measurements in synthetic dSTORM data of known ring size by respective deep learning models. Underlying ground truth ring diameter is given on the left. Averaged recognized ring images are shown for each model and diameter, average ring diameter and deviation from ground truth are shown below the images. Number of rings averaged is shown on the bottom left of each image. Scale bar: 200 nm. **(C)** Deviation from respective ground truth ring diameter in percentage points measured for all models and for all 7 ground truth ring diameters shown in (B). Black lines indicate mean deviation. **(D)** Deviation from respective ground truth ring diameter in percentage plotted against ground truth diameter for all models. Dashed line marks 0% deviation, colored lines show a linear fit.

**FIGURE 6:**
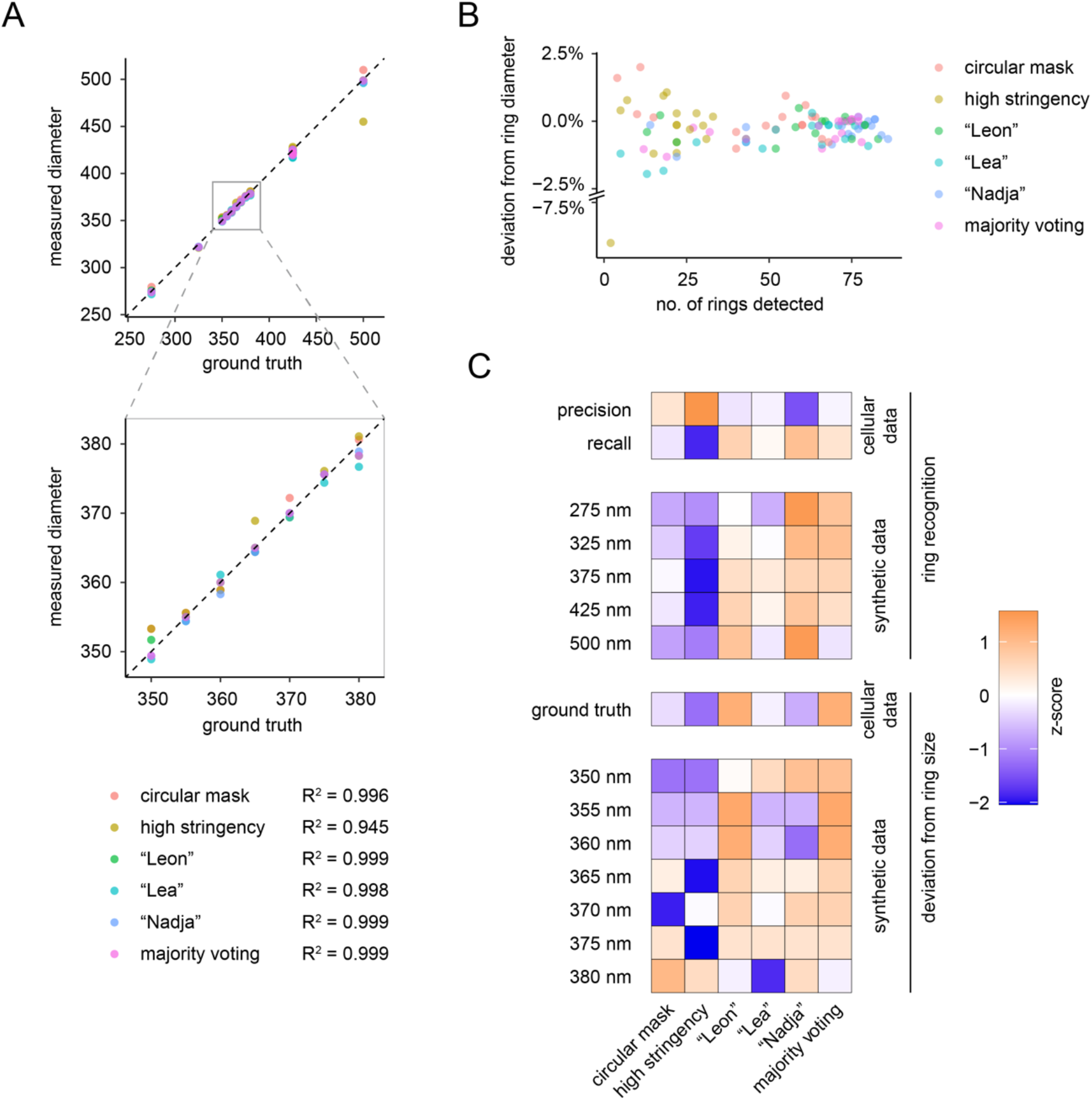
Deep learning models accurately and robustly measure septin ring size in dSTORM data. **(A)** Measured ring diameter in synthetic dSTORM data plotted against ground truth ring diameter for all models used in this study. The magnified inset below shows the ring size range exemplified in Figure 5B. **(B)** Deviation from respective ground truth ring diameter for synthetic rings from 275 nm to 500 nm in percentage points measured for all models plotted against the respective number of detected rings in each case. Note that deviation from ground truth decreases with larger number of rings detected. **(C)** Global analysis of deep learning model performance. From top to bottom: Precision and recall values in cellular data, number of rings recognized in synthetic data, ring size deviation from ground truth in cellular data and deviation from ground truth ring size in synthetic data. Heat map shows horizontally z-scored values.

It seems that the z-scores for recall and precision, respectively, are correlated with specific outcomes. The model with the lowest precision, “Nadja”, performed best in recognizing rings of sizes significantly different from those measured in native cells, suggesting that a less stringent selection of cellular structures as rings allows the model to recognize rings of different sizes more easily. On the other hand, the model with the highest precision, “high stringency”, did not perform well in synthetic data of ring sizes. For recall on the other hand, the models that most reliably recognized and most accurately measured rings in the expected size in cells, showed the highest values (“Leon” and “majority voting”), whereas the models with low recall (“high stringency” and “circular mask”) exhibited the worst performance. We concluded that precision and recall are important indicators of model performance, but they do not suffice to predict experimental outcome.

Having found models that can recognize septin rings of different sizes and accurately measure their diameter, we decided to test our models on an experimental system, where septin ring size is known to vary. Septin complexes in mammalian cells are non-polar, palindromic rod-shaped heterooctamers of around 34 nm length^40–44^ that bear two Septin-9 subunits in their middle. When Septin-9 is removed from cells through RNA interference, septin rings are known to be smaller^37^. This suggests that ring size is connected to complex length as the septin complexes created in absence of Septin-9 are known to be hexamers^45^ of around 26 nm length. If that is the case, septin ring size should reflect complex size and ring size should be dependent on the number of complexes in the ring. To ask, whether we can thus estimate the number of septin complexes in rings from SMLM measurements, we performed SMLM on embryonic fibroblast cells from *wt* and Septin-9^−/−^ mice^46^. We found septin rings to be in the size range of 300 - 400 nm and decided to analyze ring diameter using the “Leon” and the “majority voting” models as they most accurately measured the average ring diameter in that range. Indeed, both models readily recognized many rings in measured SMLM data (Figure 7A). In *wt* cells, we measured an average ring diameter of 413.3 nm (“majority voting”) or 425.5 nm (“Leon”). Geometrically, this would translate into 39 septin octamers with a 9.2° association angle (“Leon”) or 38 octamers with a 9.4° association angle (“majority voting”). On the other hand, in Septin-9^−/−^ cells, we measured ring diameters of 334.4 nm (“Leon”) or 335.0 nm (“majority voting”), which would translate into 40 septin hexamers with an 8.9° association angle for both models (Figure 7B). Strikingly, we could thus determine that septin ring size in our cells must be determined by the propagation angle of about 9° between polymerizing septin complexes that would then lead to 40 septin complexes required to form a ring. We concluded that machine learning supported SMLM superresolution microscopy could reveal information on the ultrastructural arrangement of septin complexes in rings.

**FIGURE 7:**
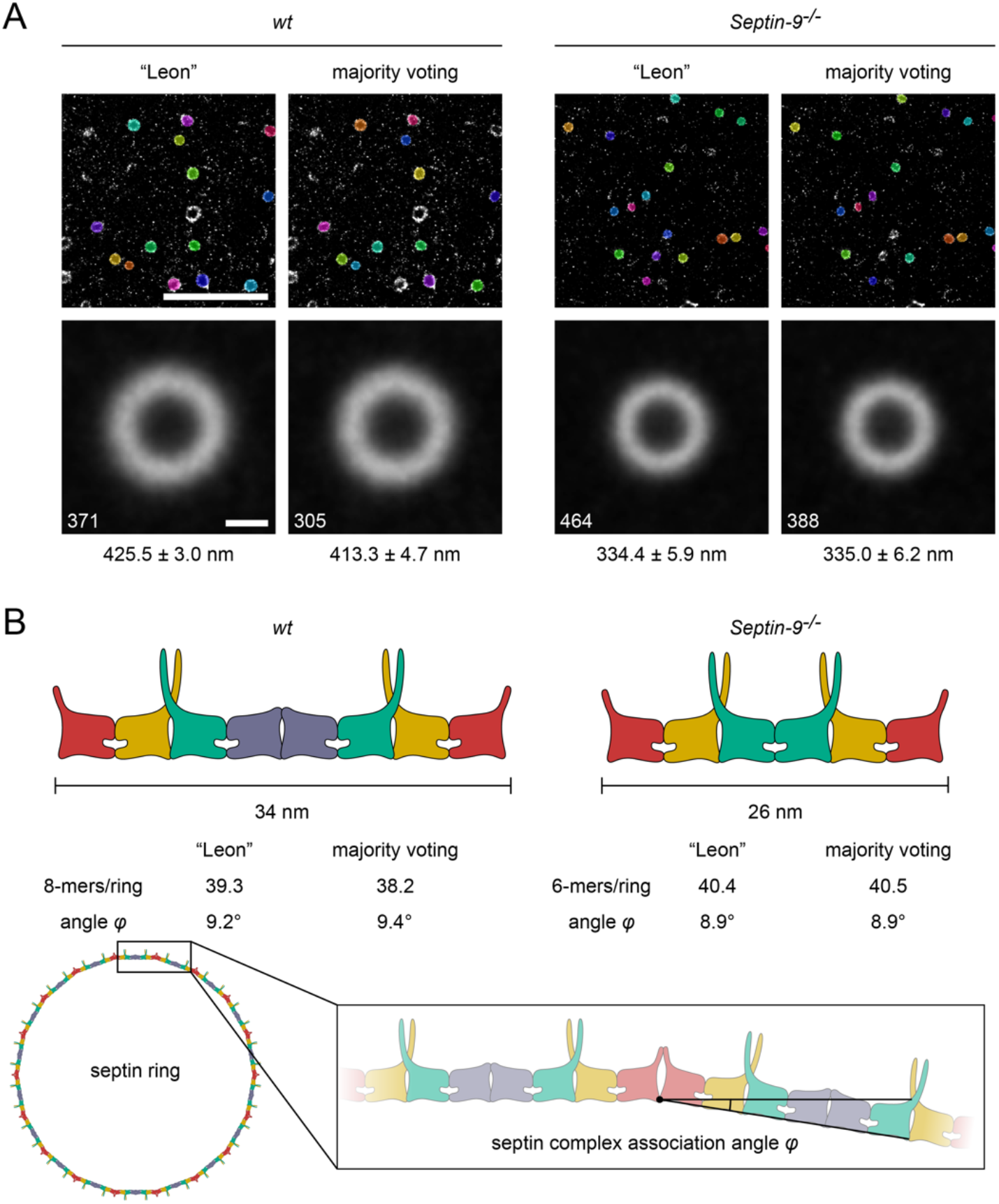
Accurate measurement of cellular septin ring size reveals the geometric architecture of septin rings. **(A)** Septin ring recognition in wild-type *(wt)* and Septin-9^−/−^ MEFs. Septin ring recognition was performed with the two deep learning models “Leon” and “majority voting”. Representative overlay of randomly colored masks of recognized septin ring structures in dSTORM images and averaged ring images are shown for each case. Average ring diameter is shown below. Number of rings averaged is shown on the bottom left of each image. Scale bars: upper row 5 μm, bottom row 200 nm. **(B)** Model of septin ring assembly. Shown are schematic representations of septin complexes. In mouse fibroblasts the knockout of Septin-9 is thought to remove the central Septin-9 dimer (blue) from the palindromic, heterooctameric complex, resulting in a heterohexamer. Since the polymerization interface for septin filament assembly is the same for hexamer and octamer, the rules for assembly of rings should be the same for both complexes and resulting rings thus smaller. Septin rings measured via deep learning-assisted superresolution microscopy data analysis in Septin-9^−/−^ cells are smaller than wildtype rings, consistent with a constant mode of assembly of rings from ~ 40 complexes with an interface angle *φ* of ^~^ 9°.

## Discussion

We here developed a deep learning-based assay for the recognition and measurement of septin ring size in superresolution images of mammalian cells. We trained deep learning models in several different ways to investigate possible bias. We demonstrate that the outline of the annotation mask, the stringency in annotation, and the generation of a consensus between experts influence the outcome of both the accuracy of measurements as well as robustness in detection of target structures. For example, we found that some of the models were more sensitive to changes in ring size than others. Especially if the models are supposed to recognize structures with a different phenotype in cells, such bias can be decisive. This suggests that benchmarking on simulated data covering the whole phenotypic range provides an excellent means to control the reliability of the deep learning models.

So far, deep learning models for the recognition of fluorescence labeled structures in cells have mostly been used to recognize structures of low or highly predictable variability such as cellular nuclei^17,38,47^. The correct recognition of different phenotypes is, however, an essential property of data analysis models in experimental biology. To ensure this is possible for our models, we use a wide array of synthetic SMLM data to test our models on the capability to recognize phenotypes that are different from what is observed in *wt* cells. Our data suggest that very stringent criteria in the selection of native rings may lead to overfitting and thus be detrimental to the recognition of rings of different size in cells, as observed for the “high stringency” and likely also the “Lea” model. On the other hand, less stringent criteria lead to a slightly lower accuracy in the measurement of ring diameter, as observed for the “Nadja” model. From all of our models, the “majority voting” model belonged to the best performing models both for recognizing *wt* cellular rings as well as for recognizing rings of significantly different size. It also resulted in very low error in ring size determination. Our data agree with previous observations that consensus or majority voting models often perform better than models based on individual annotators^48^.

Our data provide insight into the requirements in experimental design of deep learning-based data analysis pipelines of superresolution microscopy specifically and into the generation of robust deep learning models in general. We furthermore provide a robust quantitative assay for the investigation of subresolution structures assembled from an essential cytoskeletal component, the septins.

Our finding that the septin ring diameter both in Septin-9 *wt* and in Septin-9^−/−^ cells can be explained by 40 complexes adding an angle of 9° each towards ring closure suggests that superresolution microscopy of septin complexes may be useful as a readout to investigate septin complex assembly in the future. That the angle in Septin-9 containing complexes is the same as in Septin-9-free complexes suggests that the 9° propagation angle is not introduced at the center of the hexamer, but rather at the polymerization interface between terminal Septin-2-group septins in adjacent complexes. In baker’s yeast *Saccharomyces cerevisiae*, it is known that the terminal septin in the heterooctamer, here *cdc11* or *shs1*, determines the degree to which septin filaments bend. Straight filaments form with *cdc11* and ring-like filaments form with *shs1* at the termini of the complex^49^. It is tempting to speculate that the Septin-2-group septins, which occupy the termini of the mammalian octamer may play a similar role. It is known that the Septin-2 group members (Septin-1, Septin-2, Septin-4, and Septin-5) can replace one another at position X in a Septin-X/6/7/7/6/X heterohexamer biochemically^50^ and that they exhibit non-overlapping functions^36,51–53^ that may be a result of their mode of filament assembly. In the future, our assay will allow for investigations into potential influences of complex-terminal septin subunits on ring-size and thus septin polymerization properties.

## Materials and methods

### Cell culture and immunostaining

The NRK52E-Septin-2-GFP^EN/EN^ genome edited cell line expressing mEGFP-Septin2 from both alleles was generated as described^54^. All steps in the following were carried out at room temperature unless otherwise stated. NRK52E-Septin-2-GFP^EN/EN^, Septin-9^c/c^ (*wt*) or Septin-9^−/−^ (knockout) mouse embryonic fibroblasts^46^ were seeded on round coverslips (thickness no. 1.5) coated with 0.01% (w/v) poly-L-Lysine solution (Sigma) and grown in DMEM (Gibco, Thermo Fisher Scientific) supplemented with 10% (v/v) fetal calf serum, 2 mM L-Glutamine (Gibco, Thermo Fisher Scientific) and 100 U/ml penicillin-streptomycin (Sigma) at 37 °C and 5% CO_2_ in a humidified incubator. The next day, cells were treated with 5 μM Cytochalasin D for 30 min at 37 °C in DMEM without supplements to induce septin ring formation. Cells were fixed in 4% (w/v) paraformaldehyde solution in PHEM buffer (60 mM PIPES-KOH pH 6.9, 25 mM HEPES, 10 mM EGTA, 1 mM MgCl_2_) for 15 min at 37°C. 1.5. Fixation was quenched with 50 mM NH_4_Cl in PHEM buffer for 7 min. Cells were permeabilized using 0.25% (v/v) TritonX-100 in PHEM buffer for 5 min and then blocked with Image-iT FX signal enhancer (Invitrogen, Thermo Fisher Scientific) for 30 min before blocking for 1h with 4% (v/v) horse serum, 1% (w/v) BSA and 0.1% (v/v) TritonX-100 in PHEM buffer. The sample was then stained with 10 μg/ml anti-GFP nanobody coupled to Alexa Fluor-647 at 4°C overnight or with rabbit anti-human-Septin-7 IgG (1:500, IBL America) overnight and subsequently with goat anti-rabbit Alexa Fluor-647 conjugate IgG-Fab (1:500, Thermo Fisher Scientific) for 45 min in PHEM buffer supplemented with 1% (w/v) BSA and 0.1% (v/v) TritonX-100. Samples were post-fixed in 4% (w/v) paraformaldehyde in PHEM buffer for 10 minutes and quenched as described above. For the cellular septin ring dataset NRK52E 13 samples were prepared out of which 6 were stained with anti-GFP nanobody and 7 were stained with anti-Septin-7 antibody resulting in 288 and 240 images, respectively. Out of these 528 images 32 were randomly selected to be used as test dataset while the remaining 496 images were used for training of the deep learning models. For the Septin-9^c/c^ or Septin-9^−/−^ mouse embryonic fibroblasts 96 images were generated from each sample.

### Spinning disc confocal microscopy

Images were acquired on an inverted IX71 microscope (Olympus) equipped with a CSU-X1 spinning disk unit (Yokogawa) and an iLas laser illumination system (Gataca Systems) with a 491 nm laser for illumination. A 60x NA 1.42 oil objective (Olympus) was used, and images were taken with an ORCA Flash 4.0LT sCMOS camera (Hamamatsu). The System was operated using the software MetaMorph.

### dSTORM of septin rings

Coverslips with immunostained samples were mounted in a mounting chamber and filled with GLOX + BME buffer (4% (w/v) glucose, 140 mM β-mercaptoethanol, 10 mM NaCl, 200 mM Tris-HCl pH 8, 500 μg/ml glucose oxidase, 40 μg/ml glucose catalase and 10% (v/v) glycerol). dSTORM images were acquired on a Vutara 352 superresolution microscope (Bruker) equipped with an ORCA Flash4.0 sCMOS camera (Hamamatsu) and a 60x NA 1.49 ApoN oil immersion total internal reflection fluorescence (TIRF) objective (Olympus), yielding a pixel size of 98 nm. Data were acquired in TIRF illumination at a laser power density of ~35 kW/cm^2^ using a 639 nm laser. dSTORM images were reconstructed from 10,000 images taken with an exposure of 20 ms per image. Reference epi-fluorescence images were taken with an exposure of 500 ms and low laser power density. Images were reconstructed with a pixel size of 10 nm using the software Vutara SRX v.6.04.14.

### Generation of synthetic dSTORM ring data

Synthetic storm data was generated using FluoSim^39^. Geometry files were created using a python script, which generates non-overlapping randomly spaced rings with a thickness of 150 nm. Each geometry file contains 100 rings of the same radius. To generate background geometry files a python script was used, which creates 100 overlapping irregular icosahedrons per geometry with random side lengths of up to 500 nm. The geometry files were imported into FluoSim. To mimic the immunostaining process 10000 dyes were distributed randomly onto the geometries. Single-molecule localization reconstructions were then generated using parameters which resemble the experimental STORM imaging parameters (pixel size 10 nm, switch on rate 0.027 Hz, switch off rate 10 Hz, simulation time step 20 ms, localization precision 10 nm, 10000 frames).

### Data preparation for deep learning

The deep learning trainings have been carried out on 496 cellular SMLM reconstructed images of size 1024 by 1024 pixels rendered at a pixel size of 10 nm, where the number of septin rings per image was ranging from 0 to ~30 rings. Each training image has been manually annotated using each of the six different annotation strategies resulting in six different paired sets of raw and corresponding mask images. The manual annotation was performed using the Fiji plugin Labkit (https://imagej.net/plugins/labkit/). This way, our annotators manually drew masks around each potential septin ring resulting in mask images in which all pixels within an annotated ring were labeled with a distinct integer, while background pixels were labeled with zero. Based on the model as well as the number of septin-like rings in the image, a few to tens of seconds are spent on manual annotation of a single image. Moreover, the “majority voting” annotation - where the certainty level of the labels is high - is created from the annotations made by the three expert annotators. A given pixel in the “majority voting” label image is considered as a part of a mask only if it has been annotated by at least two individual annotators, otherwise it is considered background. The unseen test dataset comprising 32 cellular SMLM images was manually annotated by a septin biologist and the result was used as ground truth for further benchmarking of the models.

### Deep learning model training and parameter settings

For septin ring recognition, we trained each of the six models from scratch with the same cellular SMLM image data paired with the corresponding set of model specific mask images. To do so, we selected the 2D variant of the StarDist^38^ machine learning network with the U-Net structure as its core. We trained the StarDist network on the ZeroCostDL4Mic platform ^13^ as it provided simple access to a range of popular deep learning networks including StarDist. The hyperparameters of the StarDist structure, originally designed for cell and nuclei segmentation problems, were tuned in our implementations for the recognition of rings in SMLM image data. To do so, we adjusted the ZeroCostDL4Mic StarDist notebook to enable a long-running grid search for hyperparameter optimization as well as offline data analysis. The grid search yielded different sets of hyperparameters when optimizing for recall and precision, of which we chose the one that ensured a reasonable balance between the two criteria in all the six trained models. Table 1 summarizes the values of the StarDist network’s hyperparameters used in our experiments. In order to avoid adding another cause for bias, no augmentation was used for training. Furthermore, transfer learning is not considered in our work. The batch size was set to 8 and a mae function was chosen as the loss function. Of the training dataset, 10% were used as a validation set to monitor possible overfitting during training.

**Table 1.** Hyperparameter configuration of the Stardist network used in this research

| Parameter | Value |
| --- | --- |
| Number of epochs | 200 |
| Patch size | 1024x1024 |
| Batch size | 8 |
| Percentage validation | 10 |
| n rays | 32 |
| Grid parameter | 2 |
| Initial learning rate | 0.0003 |
| Augmentation | No |
| Transfer learning | No |

To guarantee a fair comparison of the six different annotation strategies, we trained each pair of raw images and corresponding mask images on the same StarDist network with fixed hyperparameters. In order to train the StarDist network, the key adopted python packages included tensorflow (v 0.1.12), Keras (v 2.3.1), csbdeep (v 0.6.3), numpy (v 1.19.5), and cuda (v 11.1.105). The training process has been accelerated using a Tesla P100 GPU and depending on the model it took between 123 to 161 minutes.

### Ring recognition by means of trained models

For benchmarking, we ran the six trained models on the cellular test dataset and the synthetic ring dataset using the ZeroCostDL4Mic platform. For the ring recognition in the Septin9^c/c^ and Septin9^−/−^ cell data we exported the models from the ZeroCostDL4Mic platform and ran them using the StarDist Fiji plugin. The outputs generated by the models were mask images of the same size of the corresponding test image.

### Ring averaging and diameter measurement

To generate average ring images, the model output mask images were converted into binary images using Fiji. The centers of the individual ring masks were calculated and fixed-size bounding boxes were created around the center points. Bounding boxes touching the image borders were excluded from further measurements. The bounding box ROIs were then together with the corresponding images fed to CellProfiler^47^ to extract per-ring crops from the corresponding raw images. Using a custom written Fiji macro, we measured the average ring diameter as follows: the ring crops per condition were averaged and the diameter was determined by measuring horizontal line profiles with a line thickness of 1 pixel on 18 successive 10° rotations of the image (see Figure 1C). In each of these 18 intensity plots the peak-to-peak distance was measured and then averaged to calculate the ring diameter.

### Calculation of septin ring geometry

The septin ring circumference *C* was calculated based on the formula *C* = *π*d* where *d* is the measured ring diameter. The circumference was then divided by the approximate periodicity length of heterooctameric or heterohexameric complexes as can be found in filaments including the Septin-2-NC interface gap in the open conformation as predicted by pdb structure data (pdb codes: 2QAG, 7M6J, 6UQQ, 5CYO). This was in line with the observed periodicity length of the heterooctameric complex as found in filamentous septin assemblies^40–44^. The septin complex association angle was calculated by dividing 360° by the number of septin complexes found in the ring.

### Statistics

We used the precision metric to show how relevant the recognized objects are in comparison to the septin rings in the ground truth. It was calculated as *TP / (TP + FP)*, where *TP* stands for the number of true positives and *FP* is the number of falsely recognized objects. Recall, on the other hand, was used to show how well the recognized objects match the septin rings in the ground truth. Thus, it was calculated as *TP / (TP + FN)*, where *FN* is the number of relevant septin rings in the ground truth that were not recognized by the models. Both measures fall in the [0%, 100%] interval, with higher values corresponding to better septin ring recognition performance. *R^2^* values were calculated in R (R Project) using the “lm” function. *z*-score values were calculated across the six models using the formula *z* = *(x-μ)/σ*, where *x* is the variable (recall, precision, ring diameter, no of rings detected), *μ* is the mean over the group of variables of all six models and *σ* is the standard deviation from the mean.

## Acknowledgements

This work was funded by the Deutsche Forschungsgemeinschaft (DFG, German Research Foundation) as part of TRR 186 (Project Number 278001972) and SFB 958 and core funding through Freie Universität Berlin. SMLM imaging was performed in the Core Facility BioSupraMol facility of Freie Universität Berlin. The authors wish to thank Richard C. Garratt for help with the measurement of the septin heterooctamer dimensions based on pdb structures. Furthermore, the authors wish to thank all members of the Ewers laboratory for helpful discussions.

